# C6orf203 controls OXPHOS function through modulation of mitochondrial protein biosynthesis

**DOI:** 10.1101/704403

**Authors:** Sara Palacios-Zambrano, Luis Vázquez-Fonseca, Cristina González-Páramos, Laura Mamblona, Laura Sánchez-Caballero, Leo Nijtmans, Rafael Garesse, Miguel Angel Fernández-Moreno

## Abstract

Mitochondria are essential organelles present in the vast majority of eukaryotic cells. Their central function is to produce cellular energy through the OXPHOS system, and functional alterations provoke so-called mitochondrial OXPHOS diseases. It is estimated that several hundred mitochondrial proteins have unknown functions. Very recently, C6orf203 was described to participate in mitochondrial transcription under induced mitochondrial DNA depletion stress conditions. Here, we describe another role for C6orf203, specifically in OXPHOS biogenesis under regular culture conditions. HEK293T *C6orf203*-Knockout (KO) cells generated by CRISPR/Cas9 genome editing showed both reduced grow in galactose, as a carbon source, and in their oxygen consumption capability, strongly suggesting an OXPHOS dysfunction. *C6orf203*-KO also provoked a depletion of OXPHOS proteins and decreased the activity of the mitochondrial respiratory chain complexes. C6orf203 was present in high molecular weight complexes compatible with mitoribosomes, and *in vivo* labelling of *de novo* mitochondrial proteins synthesis revealed that *C6orf203*-KO severely but not completely affected the translation of mitochondrial mRNAs. Taken together, we describe herein a new function for C6orf203, making it a potential OXPHOS disease-related candidate.

## Introduction

Mitochondria are characteristic double-membrane organelles, consisting of an outer mitochondrial membrane (OMM) and an inner mitochondrial membrane (IMM). The space between both membranes is known as the intermembrane space and that surrounded by the IMM is known as the mitochondrial matrix. The IMM forms numerous folds known as the mitochondrial cristae. Their central function is the production of most of cellular energy (in the form of ATP) through coupling ADP phosphorylation to the dissipation of a proton gradient generated by proton pumping from the mitochondrial matrix to the intermembrane space. This process is executed by the oxidative phosphorylation (OXPHOS) system, formed by the four complexes (I to IV) of the electron transport chain plus the ATP synthase or Complex V, which is located at the cristae [1]. Nevertheless, mitochondria are involved in other important cell physiological processes such as fatty acids β-oxidation, pyrimidine synthesis, iron sulfur clusters-key cofactors synthesis [2, 3], apoptosis [4], calcium homeostasis [5], reactive oxygen species (ROS) production [6] or cAMP/protein kinase A (PKA) signalling [7]. The mitochondria house their own genome, the mtDNA. The mammalian mtDNA is a 16,569 bp double-strand DNA (dsDNA) molecule which is organized and maintained within the nucleoids located in the mitochondrial matrix [8]. In mammals, the mtDNA encodes for 13 proteins (all of which are subunits of the OXPHOS system), 2 ribosomal RNA (rRNA) and 22 transfer RNA (tRNA) [9]. In the past years, novel open reading frames have been described in the mtDNA, such as humanin [10], the mitochondrial open reading frame of the 12S rRNA-c (MOTS-c) [11] and the small humanin-like peptide (SHLPs) [12]. In humans, it has been calculated that the mitochondrial proteome is composed of 1100-1900 proteins [13, 14]. As a consequence, ~ 99% of the mitochondrial proteins are encoded by the nuclear DNA (nDNA). These nDNA genes are transcribed in the nucleus, translated in the cytosol and subsequently the proteins are imported into the mitochondria. Defects in genes participating in the biogenesis of the OXPHOS system are often the cause of diseases. They could be due to mutations in either nDNA genes or mtDNA genes, affecting structural subunits of the OXPHOS complexes, assembly factors or mtDNA-maintaining and -decoding machineries including replication, transcription, mtRNAs processing and translation [15]. Although the human mitochondrial diseases are considered as rare diseases, their global prevalence, taken all together, is ~1 case in 8,000 in adults and ~1 case in 25,000 in children [15]. Clinically, they are a heterogeneous group of diseases, which may be revealed at any age and affecting any tissue or can be multisystemic, although they are principally linked to cells with a high energy demand. Unfortunately, the coenzyme Q_10_ (CoQ_10_) deficiency syndrome is the only group of mitochondrial diseases with an effective treatment ([16]). The identification of the specific genetic defect in a patient suspected to have a mitochondrial disease is complicated due to the heterogeneous phenotypes caused by mutations in a same gene [15]. In addition, at least the 20% of the mitochondrial proteome is uncharacterized, and the biochemical function of these proteins remains unknown ([17]). Thus, the identification of new mitochondrial genes and the molecular process that they are involved in is essential in order to completely understand the genetic basis of the mitochondrial diseases. From 2006, there have been several milestones to predict non-described genes involved in mitochondrial OXPHOS diseases such as the computer program Maestro [18]and MitoCarta [19] and its updates MitoCarta2.0 [13] and MitoCarta+ [17]. At the same time, a similar mitochondrial proteome database, MitoMiner v3.1, was developed [14] and recently updated to MitoMiner v4.0 [20]. High-throughput screening (HTS) and next generation sequence (NGS) has emerged as powerful tools to completely characterize the mitochondrial proteome. Our group has previous experience in identifying novel mitochondrial genes. We demonstrated that the coiled coil domain-containing protein 56 (CCDC56) is a mitochondrial protein essential for cytochrome *c* oxidase function in *Drosophila melanogaster* [21] and its human ortholog, COA3, stabilizes cytochrome *c* oxidase I and promotes cytochrome *c* oxidase assembly [22]. In addition, we observed that GATC was a new *Drosophila melanogaster* protein, ortholog to the corresponding subunit of the bacterial Glutamyl-tRNAGln amidotransferase, and candidate for mitochondrial localization. We described that this amidotransferase activity is essential for mammalian mitochondrial translation *in vivo* [23] and recently that mutations in any of its subunits (GATA, GATB and GATC) provoke severe mitochondrial diseases [24]. *Ccdc56* and *GatC* are transcribed in bicistronic mRNAs in *Drosophila melanogaster* together with the mitochondrial translation factor *mt-Tfb1* gene and the mitochondrial DNA polymerase *PolγB* subunit respectively [21] (Peña et al. manuscript in preparation).

The *C4884 Drosophila* gene is also transcribed in a bicistronic mRNA together with the uncharacterized gene *CG42487*. The *C4884* ortholog gene in humans is the *C6orf203* gene. The nDNA-encoded *C6orf203* gene is located in the long arm of the chromosome 6 (6q21), and it is predicted to possesses a conserved domain that belongs to the RNA-binding S4 domain superfamily (IPR036986). Interestingly, C6orf203 has been included into the MitoCarta+ and MitoMiner v4.0 databases as a mitochondrial uncharacterized protein, showing, in addition, a high score for mitochondrial location using the algorithm for prediction of mitochondrial targeting sequences MITOPROT (https://ihg.gsf.de/ihg/mitoprot.html; [25]). Moreover, previous evidences strongly suggest mitochondrial location of C6orf203. In 2011, C6orf203 was identified as a mitochondrial phosphoprotein with unknown function in human muscle [26] and it was present, together with a number of mitochondrial ribosome proteins, when the mitoribosomal subunit ICT1 was immunoprecipitated [27]. A few weeks ago, it was described that C6orf203/MTRES1 is a mitochondrial protein involved, acting at the transcriptional level, in maintaining mitochondrial RNA levels under stress mediated by mtDNA depletion provoked by BrEt treatment [28].

In this work, we describe that C6orf203 is part of the mitochondrial proteome that is evolutionarily conserved from insects and worms to mammals. We demonstrate by CRISPR/Cas9 genomic editing its participation in OXPHOS biogenesis under normal culture conditions. Moreover, we demonstrate the presence of C6orf203 in very high molecular weight complexes and its involvement in mitochondrial protein biosynthesis. We propose C6orf203 as a strong candidate to be considered in the search of genes responsible for mitochondrial pathology not necessarily severe.

## RESULTS

### C6orf203 is evolutionary conserved and contains an RNA binding domain

Previously, our laboratory showed that orthologs of some highly conserved and essential genes for the OXPHOS system biogenesis in humans such as *mt-Tfb1*, *GatC*, *Pol*γ*-*β and *Coa3* were encoded on bicistronic mRNAs in *Drosophila* [21] (Peña et al. manuscript in preparation). Since this arrangement, unusual in eukaryotes, could suggest a tendency for mitochondrial genes in this organism we delved into the search and identified the *CG4884* gene also encoded on a bicistronic mRNA. The human ortholog of *CG4884* is *C6orf203*, recently described as a mitochondrial RNA binding protein.

Although three *C6orf203* mRNA isoforms are shown (ENSG00000130349), they code only for two putative inmature proteins of 240 and 245 amino acids in length respectively. However, both of them render a unique mature functional protein of 156 amino acids according to the predicted mitochondria targeting sequence which extends amino acids 1-84 (Fig 1). C6orf203 is a protein well conserved in Metazoa, from *Drosophila* to humans, being absent in bacteria, fungi and plants. Based on Interpro [41] and NCBI CDD [42] domain annotation tools and previous work [28] C6orf203 harbors an RNA binding domain (Fig 1) essential for ribosome function in bacteria [43] and for mitochondrial transcription in humans [28].

**Fig 1.**
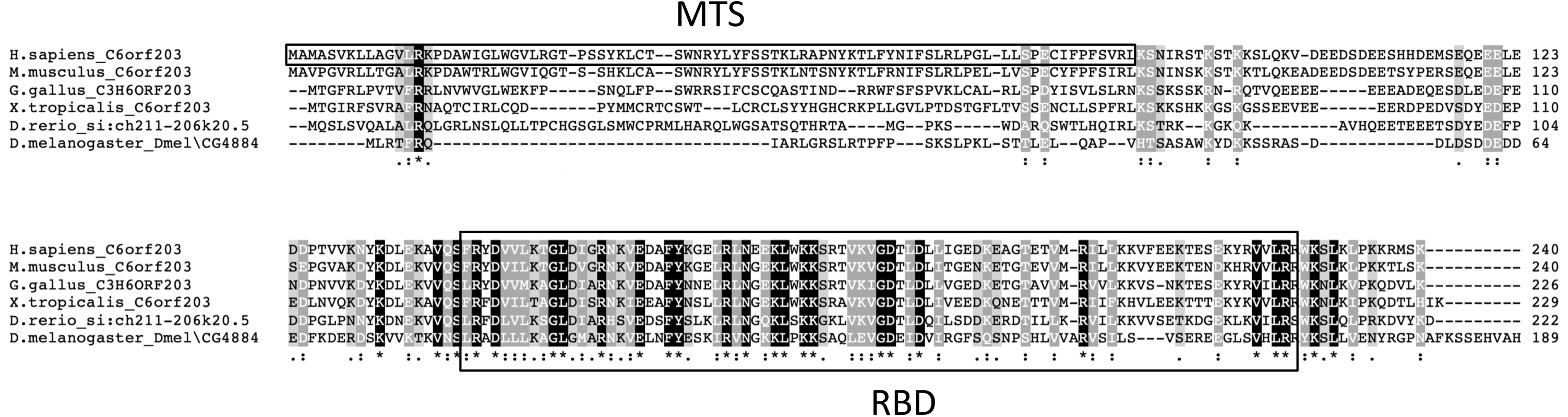
C6orf203 is an evolutionary conserved protein harboring an RNA binding domain. Alignment of human C6orf203 sequence with some orthologs was performed using Clustal Omega. UniprotKB accession numbers of sequences are: Q9P0P8 (*H.sapiens*), Q9CQF4 (*M.musculus*), F1NMS1 (*G*.gallus), F7AXE1 (*X.tropicalis*), B7ZD93 (*D.reri*o), Q9VAX9 (*D.melanogaster*). Asterik (*) under white letter over black indicates completely conserved residues, while colon (:) under white letter over grey indicates similar aminoacid properties and colon (.) under black letter on clear grey indicates minor homology. Human sequence squared named MTS shows the mitochondrial targeting sequence. The RNA binding domain (RBD), according to NCBI Conserved Domain Search [42], extends from positions 141 to 227, is squared.

### C6orf203 is a mitochondrial matrix protein that interacts with the Mitochondrial Inner Membrane

To confirm the mitochondrial localization of human C6orf203, we transiently expressed the C-terminal GFP-tagged construct in both HEK293T and HeLa cells. As shown in (Fig 2A), C6orf203-GFP colocalized with the mitochondrial marker Mitotracker Red in both cell types.

**Fig 2.**
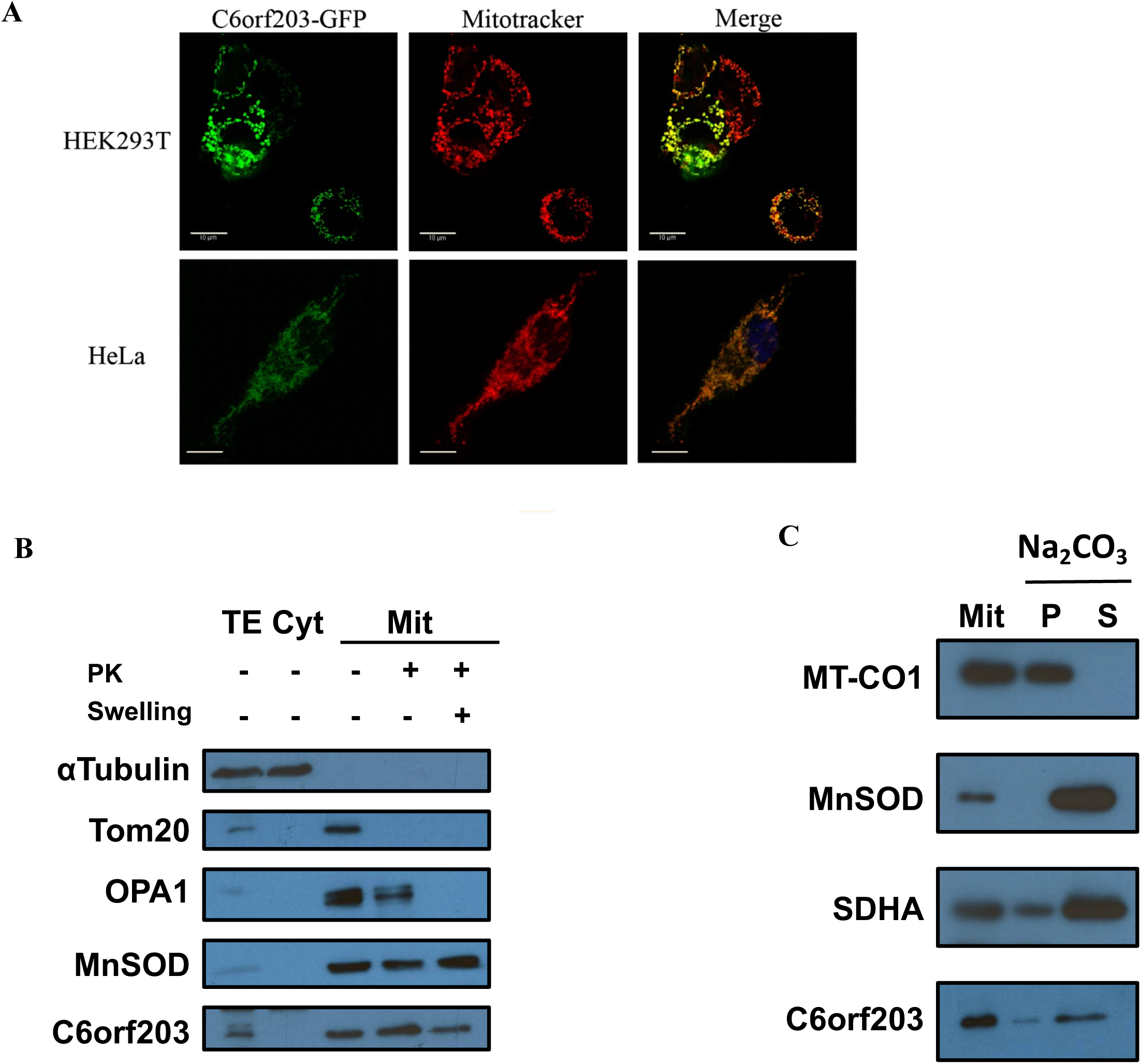
C6orf203 is located in the mitochondrial matrix. **A** Fluorescence microscopy of HEK293T and HeLa cells expressing C6orf203-GFP. Cells were transfected with plasmid peGFP-N1 expressing the C-terminal GFP-tagged C6orf203 (in green). Mitochondria were stained using Mitotracker Red (in red). Images were obtained in Leica SP5 confocal microscope. Scale bars represent 10 µm. **B** Immunoblot analysis of purified mitochondria from HEK293T after Proteinase K (PK) accessibility test. Mitochondria were left untreated or swollen to rupture the outer membrane before the treatment with PK. TE, Total cell extract. Cyt, cytosolic fraction. Mit, purified mitochondria. α-tubulin is a cytosolic marker, TOM20 is an OMM marker, OPA1 is a mitochondrial intermembrane space marker and MnSOD in a mitochondria matrix marker. **C** Immunoblot analysis of mitochondrial extract after alkaline sodium carbonate (Na_2_CO_3_) pH11 extraction. Mit, purified mitochondria. P, pellet (membrane proteins). S, supernatant (soluble proteins). MT-CO1, SDHA and MnSOD mark integral IMM, peripheral IMM and soluble matrix proteins, respectively.

To determine the sub-mitochondrial location of the endogenous C6orf203, mitochondria from HEK293T cells were isolated and incubated with proteinase K (PK) together with an iso-osmotic buffer or a hypo-osmotic buffer in order to induce mitochondrial swelling. The mitochondrial swelling causes the outer membrane rupture. Our results confirm that C6orf203 is protected from degradation by PK when mitochondria were swollen (Fig 2B), similar to MnSOD, a well-characterized mitochondrial matrix protein, strongly suggesting that C6orf203 is located within the mitochondrial matrix. In order to determine a putative association of C6orf203 to the IMM, we perform a carbonate extraction assay (Fig 2C). We used well-known selected markers such as MT-CO1 (an integral IMM protein), SDHA (a peripheral IMM protein) and MnSOD (a soluble matrix protein). The behavior of C6orf203 was similar to that of SDHA, strongly suggesting that C6orf203 is a mitochondrial matrix protein with a weak association to the internal face of the IMM as has been observed for other proteins involved in different processes such as COQ8B (coenzyme Q biosynthesis) [44] or some mitochondrial translational activators such as Mba1 [45] or PET111 [46].

In this analysis, we also observed that the molecular weight of the mature form of C6orf203 is 19 kDa, which corresponds well with the predicted N-terminal cleavage of 84 aminoacids (Mitoprot).

### Generation of *C6orf203*-KO cell lines

To obtain a human cell model to study the function of C6orf203, we generated HEK293T Knockout cells (*C6orf203-*KO) using the genomic edition system CRISPR/Cas9n(D10A) nickase as described in Materials and Methods. This enzyme was targeted by two gRNAs with specific recognition sites on different genomic DNA strands of C6orf203 exon 3 (Fig 3A). gRNAs recognition sites were separated by 76 bp, producing nicks that are recognized by the cell as a single injury creating insertions and deletions during repair [30]. The region between gRNAs encompassed the first coding exon, which includes the beginning of its RNA binding domain (Fig 3A).

**Fig 3.**
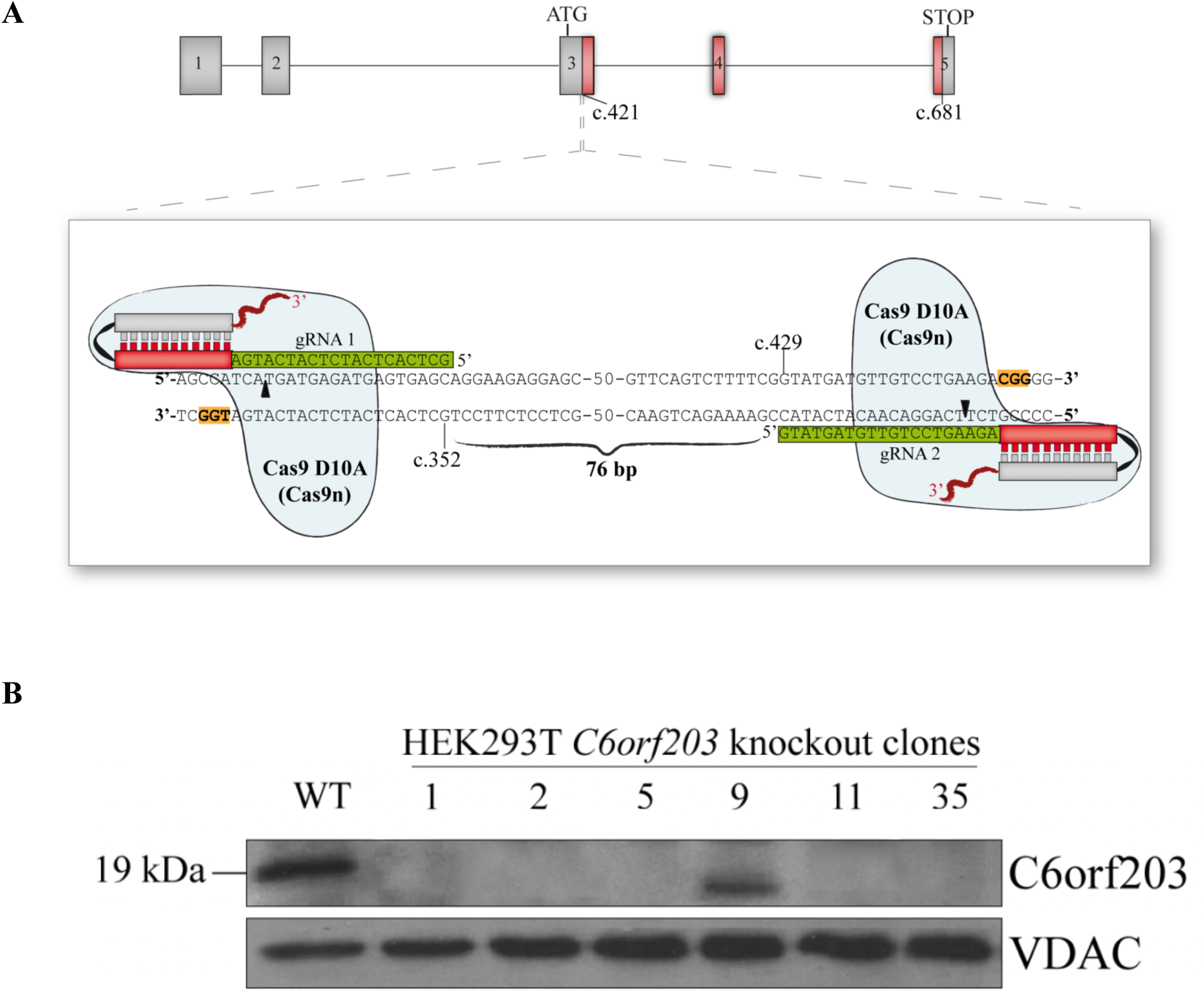
Generation of C6orf203-KO cell lines. **A** Schematic illustration of Cas9n(D10A) nickase double nicking the human *C6orf203* locus using a pair of gRNAs. Exons are shown as boxes and numbered. Coding region corresponds to the exons sequence between ATG and STOP signal. In red is shown the coding region of S4 RNA binding Domain. gRNAs, with CRISPR sequences in green, were designed to recognize the first nucleotides coding the conserved RNA binding domain showed in Fig 1. The PAM sequence is shown highlighted in orange. **B** Immunoblot analysis of mitochondrial extracts from wild type HEK293T cells (WT) and *C6orf203*-KO clones (C1, C2, C5, C9, C11, C35) using anti-C6orf203 antibody. VDAC was used as loading control. 25 μg of purified mitochondrial protein were resolved in a 12% by SDS-PAGE. All but clone 9 showed no expression of C6orf203. Clone 9 expressed a truncated form of the protein.

24 hours after transfection of HEK293T cells with plasmid expressing gRNAs and CRISPR/Cas9n(D10A) nickase, we analyzed by PCR the edited region in pooled cells as described in Materials and Methods and Fig EV 1 legend. The efficiency of edition was approximately 65%, prompting us to generate clones from the pool (Fig EV 1).

Analysis of 36 individual clones by western blot revealed that 6 of them had no detectable levels of C6orf203 (Fig 3B). The genomic analysis of these *C6orf203*-KO clones was carried out by PCR of the edited region, cloning on an *E.coli* vector and sequence 21 clones, showing deletions in *C6orf203* alleles provoking frameshifts, and therefore premature stop codons and truncated versions of C6orf203, and deletions in the final version of C6orf203 (Table EV 1).

### The absence of C6orf203 leads to a decrease in mitochondrial function

As a first approach to determine whether *C6orf203* disruption could affect mitochondrial function, we first analyzed the capability of the *C6orf203*-KO clones to growth in a medium containing galactose as a carbon source. Galactose entry on glycolysis involve several limiting enzymatic steps (Leloire pathway) [47] so that a functional OXPHOS system can maintain energetic metabolism in those conditions but a non-functional OXPHOS system cannot, severely affecting cell growing or even provoking cell death. *C6orf203*-KO cells grew slightly slower than HEK293T wild type cells in high glucose medium but were unable to grow in galactose containing medium (Fig 4A), indicating that C6orf203 is required for a proper mitochondrial OXPHOS function.

**Fig 4.**
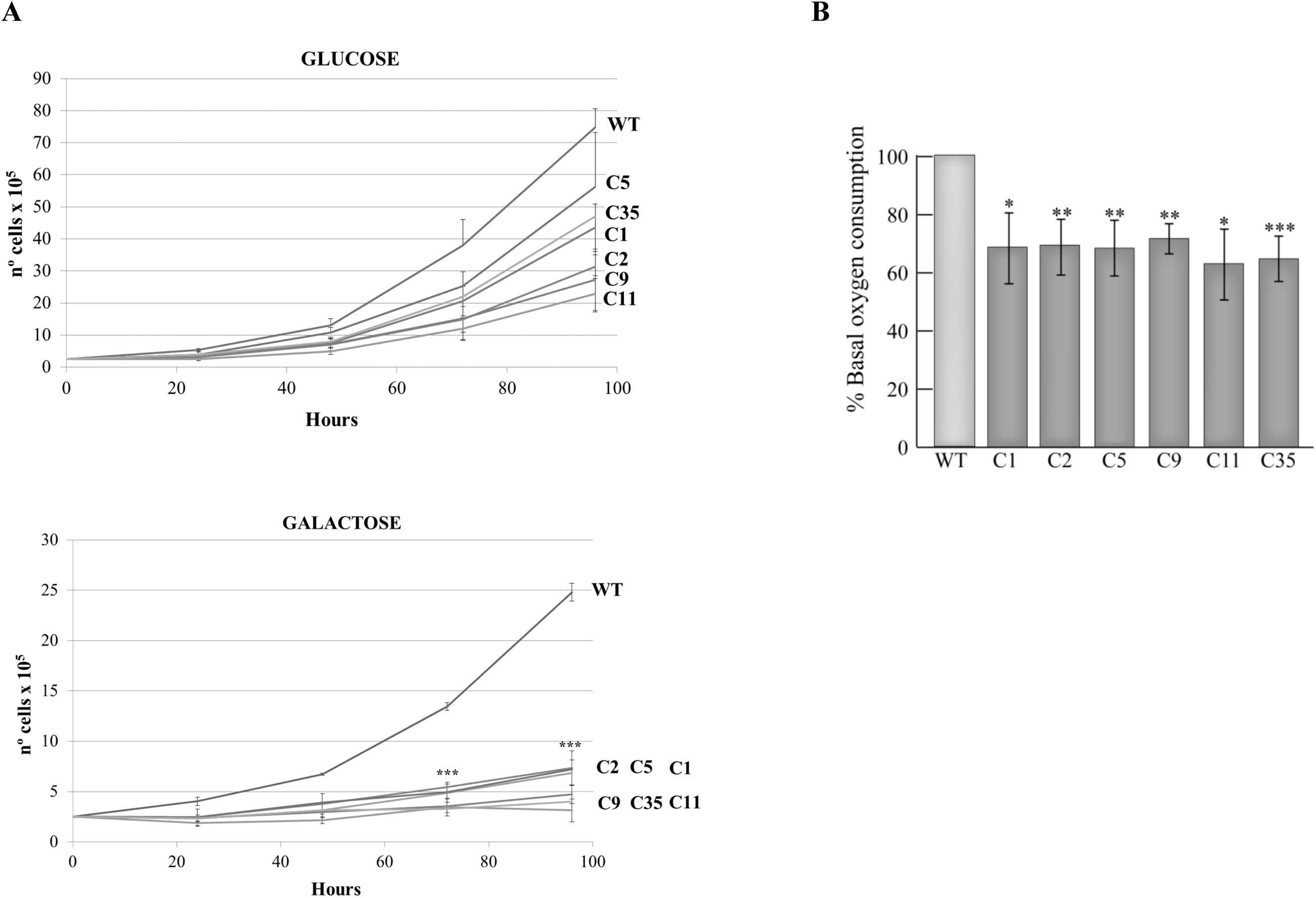
*C6orf203* knockout results in a defect in mitochondrial function. **A** Cell growth analysis of wild type cells and *C6orf203*-KO clones. Cell lines were grown for 96 hours in 4.5 g/l glucose or 4.5 g/l galactose medium to assess their mitochondrial function. Lines represent the mean of three independent experiments. Error bars represent standard deviation (SD). P values (***p<0.001 for *C6orf203* KO clones versus wild type cells) were determined by two-tailed unpaired student’s t-test. n=3. **B** Oxygen consumption of 4×10^6^ cells was measured for 30 minutes using a Clark-type O_2_ electrode in medium containing 0.9 g/L galactose. Data are represented as percentage relative to wild type HEK293T cells. The bars represent the mean of 3 independent experiments ± standard deviation. P values (*p<0.05, **p<0.01, ***p<0.001 for *C6orf203*-KO clones versus wild type cells) were determined by two-tailed unpaired student’s t-test.

In order to confirm a mitochondrial dysfunction, we next measure the oxygen consumption in intact cells. The results revealed a reduction of approximately 35% in basal oxygen consumption in the six selected *C6orf203*-KO clones when compared to control cells (Fig 4b), confirming the respiratory chain defect of cells lacking C6orf203.

For further studies, *C6orf203*-KO clones 5 (from hereon C5) and 35 (from hereon C35) were used. Thus, in order to demonstrate that the functional defects observed in *C6orf203*-KO cells were due to the absence of C6orf203, we carried out rescue studies via ectopic stable expression of the Flag-tagged version of C6orf203 in C5 and in C35 clones, rendering C5-Flag and C35-Flag cell lines (Fig 5A). C6orf203-Flag expression was able to fully rescue the growth rate defect in galactose (Fig 5B) as well as the respiration defect (Fig 5c) in both KO cell lines. This confirmed that the phenotypes observed in knockout cells was due to *C6orf203* edition, discarding an off-target effect and confirming that C6orf203-Flag was a functional version of C6orf203.

**Fig 5.**
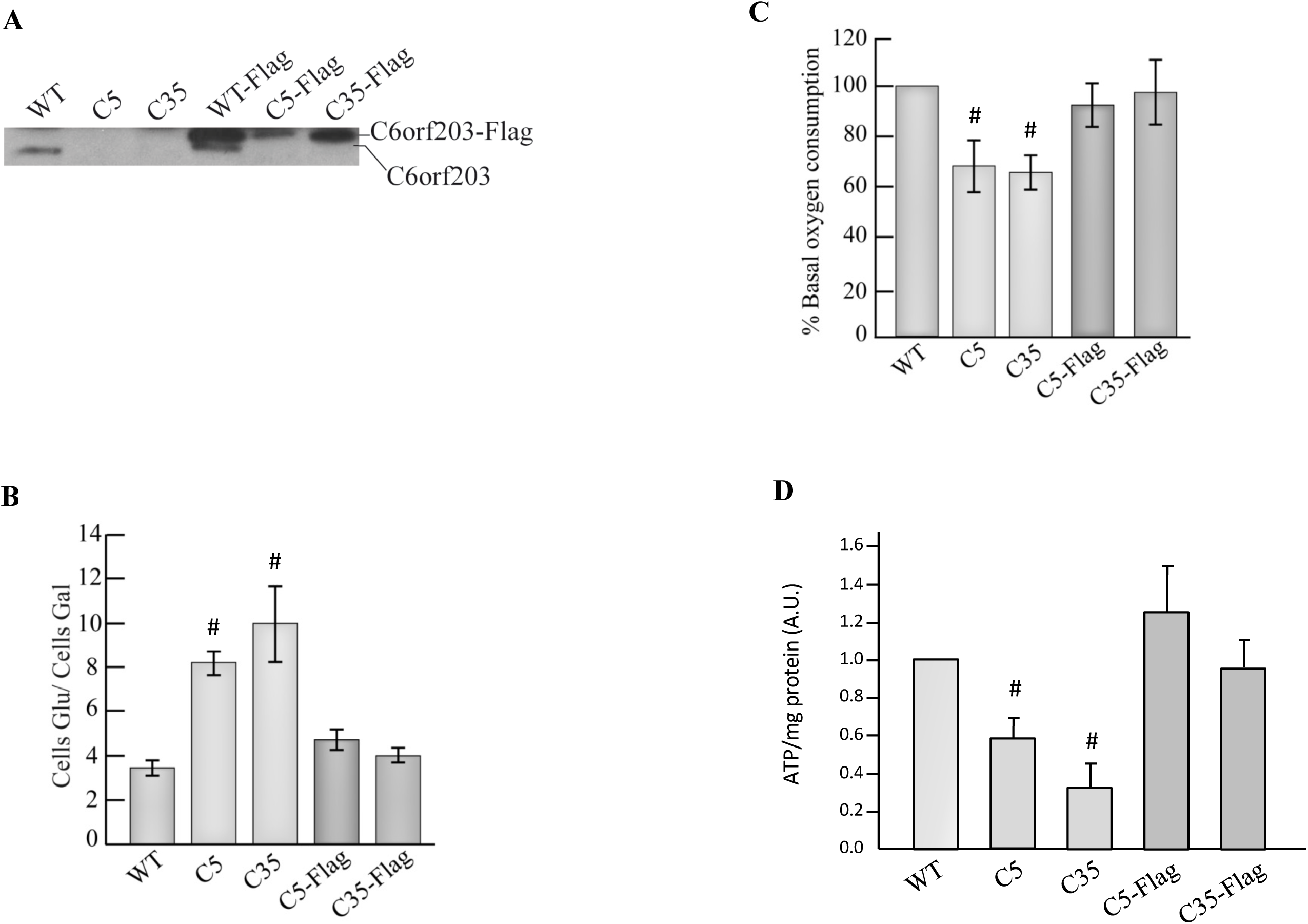
The C-terminal Flag-tagged version of C6orf203 rescue the mitochondrial defects of *C6orf203* knockout. **A** Immunoblot analysis of whole cell extracts using anti-C6orf203 antibody in wild type cells (WT), *C6orf203*-KO clones 5 (C5) and 35 (C35) and in the corresponding expressing C6orf203-Flag derivatives (WT-Flag, C5-Flag, C35-Flag). **B** Cell growth analysis of wild type cells (WT), C5, C35, C5-Flag and C35-Flag cell lines. Cell lines were grown for 96 hours in 4.5 g/l glucose or 4.5 g/l galactose medium, harvested and counted. Data are represented as the number of cells growing in glucose versus those growing in galactose. Bars represent the mean ± SD. P values (#p<0.05 for *C6orf203*-KO clones versus C6orf203-Flag transfected clones) were determined by two-tailed unpaired student’s t-test. n=3. **C** Oxygen consumption was recorded using a Clark-type O_2_ electrode in HEK293T and its derivatives C5, C35, C5-Flag and C35-Flag clones. Bars represent the mean± SD. P values (#p<0.05 for *C6orf203*-KO clones versus their corresponding C6orf203-Flag expressing clones) were determined by two-tailed unpaired student’s t-test. n= 4. **D** Mitochondrial ATP synthesis in HEK293T and its derivatives C5, C35, C5-Flag and C35-Flag clones. Cells were grown with 2-deoxy-D-glucose/1 mM pyruvate. Bars represent the mean± SD. P values (#p<0.05 for *C6orf203*-KO clones versus their corresponding C6orf203-Flag expressing clones) were determined by two-tailed unpaired student’s t-test. n= 4.

Finally, we analyzed the mitochondrial ATP synthesis under glycolysis inhibition conditions by action of 2-deoxy-D-glucose/1 mM pyruvate. Results revealed a decrease between 60%-40% in mitochondrial ATP production in C5 and C35 cell lines respectively, which reach normal values when expressed C6orf203-Flag in C5-Flag and C35-Flag cells.

In addition to the *in vivo* analysis of the carbon source dependent growing rate, the oxygen consumption capability and the mitochondrial ATP synthesis of C6orf203-KO cells compared to control cells we further investigated several other parameters of the molecular phenotype induced by lacking C6orf203 using *in vitro* approaches.

### Lacking C6orf203 leads to a reduction in the respiratory chain complexes activities

To further investigate the function of C6orf203 we analyzed if the respiratory defect observed in *C6orf203*-KO cells was the result of a defect in the respiratory chain complexes activity. Thus, we observed a reduction of 60% in complex I activity and a reduction of 50% in complex III and IV compared to control cells (Fig 6A). Activity of complex II, which is exclusively formed by nuclear encoded subunits, as well as citrate synthase, remained unchanged (Fig 6A and 6B respectively), strongly suggesting an OXPHOS alteration of mitochondrial origin. The defect in the respiratory chain complexes activity was restored by expression of C6orf203-Flag.

**Fig 6.**
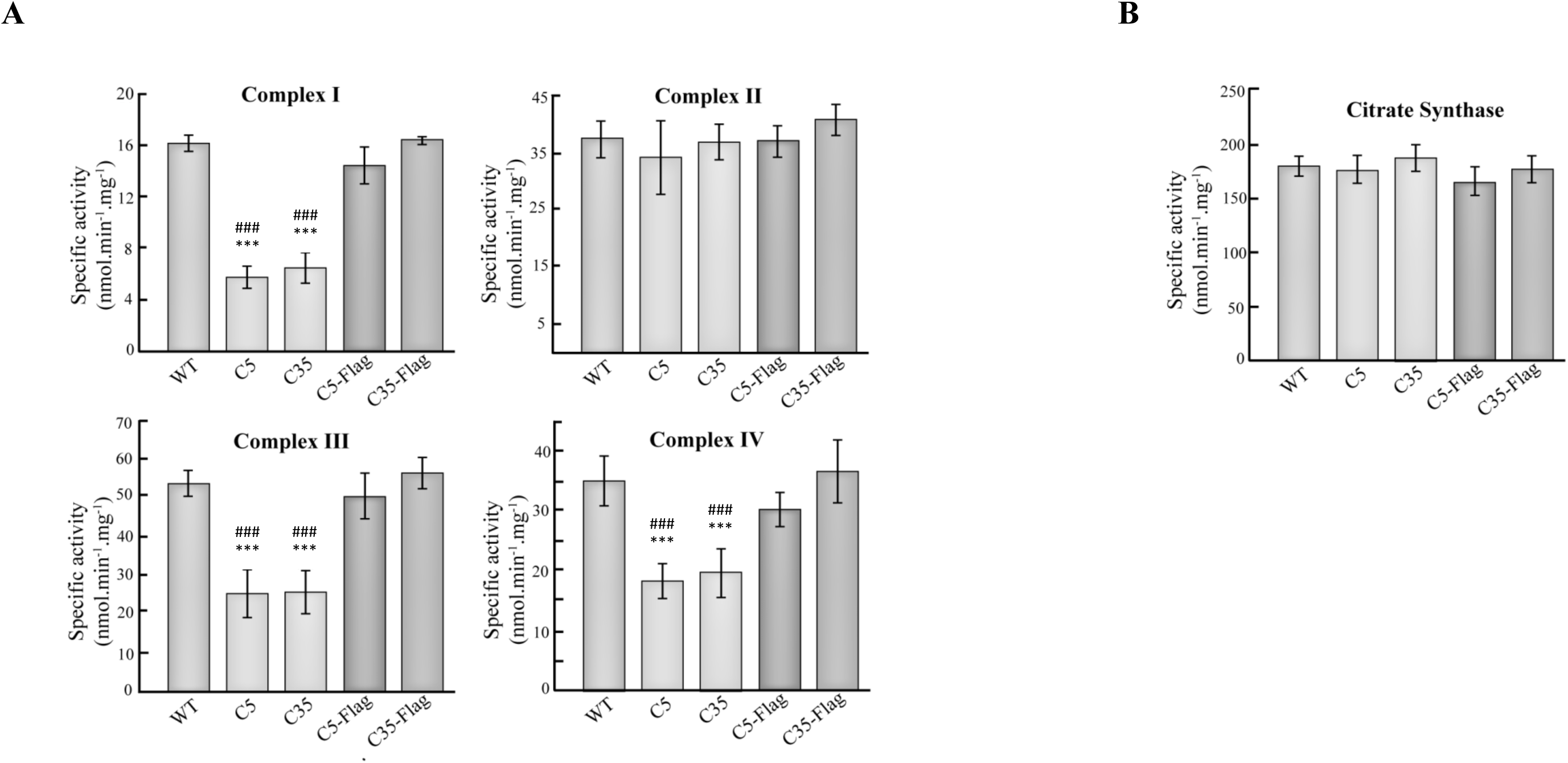
*C6orf203* knockout cells show a reduction in the respiratory chain complexes activities. **A** Specific activity of complexes I, II, III and IV were measured in control cells (WT), *C6orf203*-KO clones (C5, C35) and clones rescued with C6orf203-Flag by spectrophotometric methods. Data are presented as means ± standard deviation for *C6orf203*-KO clones versus wild type cells. ###p<0.001 for *C6orf203*-KO clones versus C6orf203-Flag transfected clones, n=4. **B** Citrate synthase activity was measured as a marker of mitochondrial mass. Data are presented as means ± standard deviation. There were no significant differences. n=4.

### C6orf203 depletion induces defects in the formation of fully-assembled OXPHOS complexes and reduces the steady-state levels of respiratory chain components

In order to gain insight regarding the reduction of respiratory chain complexes activities, we next analyzed the steady state levels of fully-assembled OXPHOS complexes. Thus, mitochondrial extracts were obtained under native conditions and resolved using blue-native PAGE (BN-PAGE). Analysis by Western Blot using antibodies against subunits of specific complexes showed a decreased in complex I (~ 60%), complex III (~ 50%), complex IV(~ 50%) and complex V (~ 50%) in *C6orf203*-KO cells compared to control cells (Fig 7a), consistent with the reduction of complexes activities seen before. Relative values of signals were measured by densitometry for each line and complex (Fig 7B). Our results also showed a reduction in fully assembled complex V and revealed accumulation of ATPase subcomplexes (Fig 7C), that were confirmed by 2D BN/SDS-PAGE (Fig 7D). These data strongly suggest that the reduction in OXPHOS complex activities is due to a decrease in the steady state levels of fully-assembled OXPHOS complexes.

**Fig 7.**
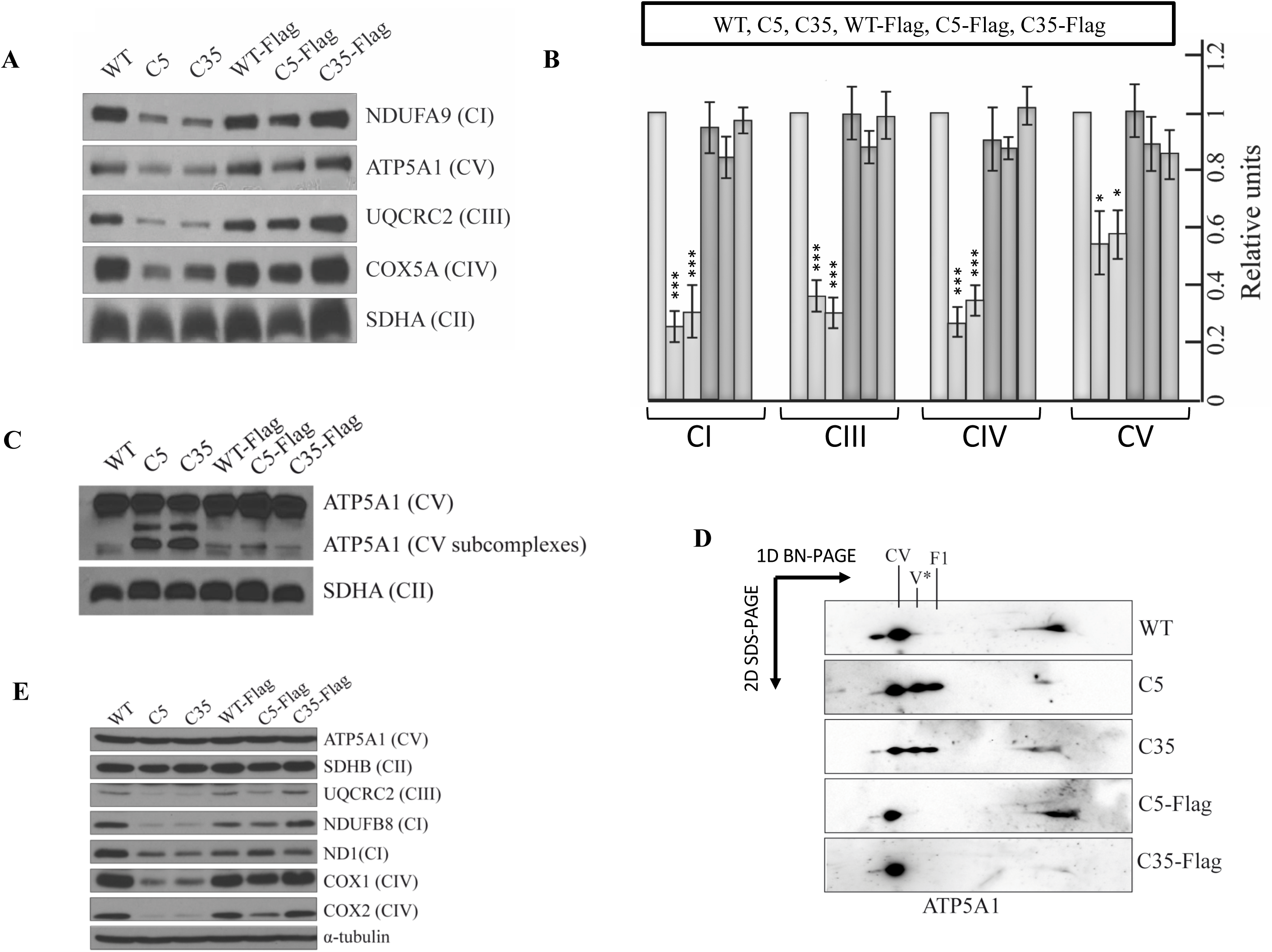
C6orf203 depletion induces defects in the formation of fully-assembled complexes and a decrease on the steady-state levels of respiratory chain components. **A** Digitonin-treated isolated mitochondria from control cells (WT), C6orf203-KO clones (C5, C35) and the C6orf203-Flag expressing clones (WT-Flag, C5-Flag and C35-Flag) were analyzed by Blue Native-PAGE. 80 μg of mitochondrial protein were resolved in 4%-15% gradient acrylamide gels and immunoblotted using antibodies against specific members of the respiratory chain complexes as indicated at the right of the panel. **B** Densitometry of signals from 4 different experiments for each cell line and each complex. Values were normalized respect to complex II that was used as loading control. The order of data presentation for each complex is shown above: WT, C5, C35, WT-Flag, C5-Flag and C35-Flag cell lines. Data are represented as means ± standard deviation of each line normalized respect control line (WT). P values (*p<0.05 and ***p<0.001 respect the control line (WT) were determined by two-tailed unpaired student’s t-test. **C** Blue Native-PAGE and immunoblot using anti-ATP5A1 antibody to analyze complex V subcomplexes in control cells (WT), C6orf203 KO clones (C5, C35) and C6orf203-Flag expressing clones. Exposure of ATP5A1 (CV) incubated membrane is higher than in panel A in order to give a better information about CV subcomplexes. **D** After Bue native-PAGE, samples were separated by a second dimension (2D BN/SDS-PAGE) and analyzed by immunoblot using anti-ATP5A1 antibody to observe complex V subcomplexes reinforcing results from C. **E** Representative immunoblot of WT, C5, C35, WT-Flag, C5-Flag and C35-Flag cell lines. 50 μg of cell protein extract were resolved by 12% SDS-PAGE. Western blot membranes were probed with the specified antibodies shown at the right of the panel. α-Tubulin was used as loading control.

We next determined whether changes in OXPHOS structural subunit steady-state levels were responsible for the alteration of assembled complexes. For that purpose, we performed western blot analysis using antibodies for a representative number of OXPHOS subunits (Fig 7E). We observed a clear reduction of the mitochondrial-encoded complex IV subunits MT-CO1 and MT-CO2 as well as MT-ND1, a complex I subunit, in *C6orf203*-KO cells compared to wild type cells. We also observed a reduction in NDUFB8 and UQCRC2 levels, two nuclear-encoded complex I and complex III subunits, respectively, which are unstable if are not incorporated into their corresponding complexes [24]. Taken together, these observations explained the sum of our previous results, and demonstrate that the absence of C6orf203 leads to a reduction of the steady state levels of mtDNA encoded OXPHOS subunits.

### *C6orf203* knockout leads to a reduction in the mitochondrial protein synthesis *de novo*

A global reduction of the levels of OXPHOS subunits may be fundamentally due to mtDNA depletion, defects in mtRNA synthesis and/or processing or dysfunctional mitochondrial protein synthesis. Thus, we first analyzed the mtDNA content of Hek293T and *C6orf203*-KO cells C5 and C35 by qPCR. No significant changes were observed (Fig EV 2), demonstrating that the OXPHOS defects associated to the lack of C6orf203 expression were not a consequence of mtDNA depletion.

Then, we analyzed the levels of mitochondrial RNAs of the same cell lines by RT-qPCR using specific probes for *12S-rRNA*, *MT-CO1, MT-CO2, MT-*ND1 and *MT-ND6* genes. As it is shown in (Fig EV 2B), no significant changes were obtained, strongly suggesting that the absence of C6orf203 had no effect on the steady-state levels of mtRNAs. In order to confirm this point and to analyze the processing of mtRNAs, we carried out northern experiments. Thus, after resolving total mtRNA from Hek293T, C5 and C35 cell lines on agarose-formaldehyde gels, we transferred them to nitrocellulose and hybridized with specific ssRNA antisense probes corresponding to *16S-rRNA* gene (or *MT-RNR2)*, *MT-CO1* and *MT-ATP8* genes. No changes were observed either in the amount or in mtRNAs intermediates (Fig EV 2C), strongly suggesting that there is no alteration of mtRNA processing in *C6orf203* knockout cells and confirming the previous evidence that the steady-state levels of mtRNAs do not change.

Finally, we performed pulse-labeling experiments to analyze the *de novo* mitochondrial protein synthesis. Thus, WT, C5, C35, C5-Flag and C35-Flag cell lines were treated with emetine to inhibit cytoplasmic translation and then ^35^S-Methionine was added to the medium for 90 minutes to incorporate specifically into the mitochondrial translation products. We found that absence of C6orf203 significantly reduces the mitochondrial capacity to translate their mt-mRNAs (Fig 8A). Densitometry of translation products and analysis estimated this reduction in approximately 35%-40% (Fig 8B).

**Fig 8.**
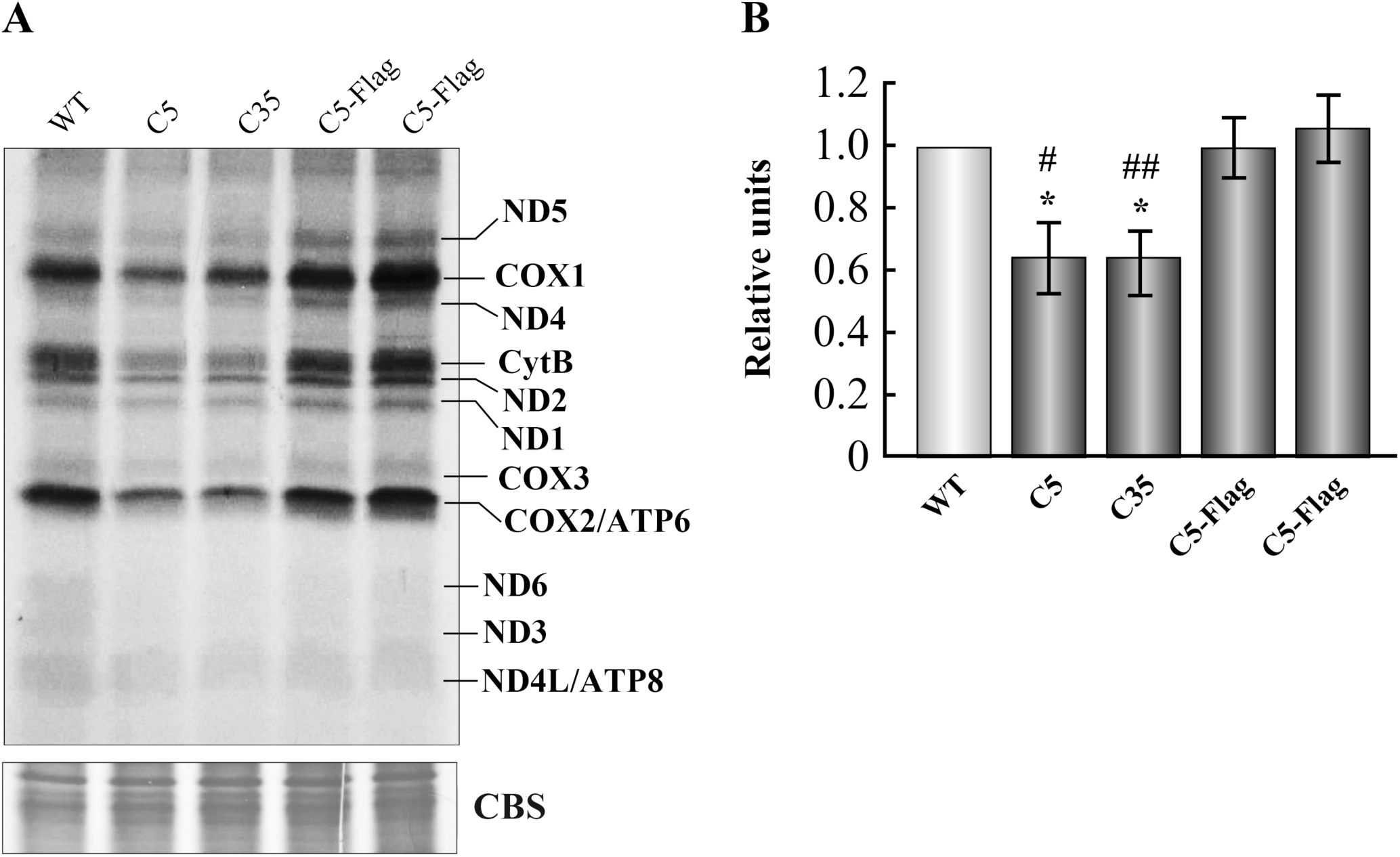
*C6orf203* depletion leads to a reduction of the mitochondrial protein synthesis *de novo*. **A** Pulse-labelling of mitochondrial translation products in control cells (WT), C6orf203-depleted clones (C5, C35) and rescued clones (C5-Flag, C35-Flag). Newly synthesized mitochondrial polypeptides were labeled with ^35^S-Methionine for 90 min. Bands corresponding to all 13 mitochondrial proteins were detected through direct autoradiography. CBS, Coomassie blue stain was used as loading control. **B** Densitometry of labelled products in panel A. Data represent mean values relative to WT cells ±SD, * p<0.05 for *C6orf203*-KO clones versus wild type cells; #p<0.05, ##p<0.01 for *C6orf203*-KO clones versus their corresponding C6orf203-Flag expressing cell lines. n=3

These data demonstrate that C6orf203 is involved in the OXPHOS system biogenesis through its participation in a very important but not essential manner in mitochondrial protein biosynthesis.

### C6orf203 forms high molecular weight complexes

Very recently, Kotrys et al. described the involvement of C6orf203 in mitochondrial transcription under induced mtDNA depletion and its interaction with mitochondrial RNA polymerase and with TFAM [28], proteins well known to be involved in the synthesis of mtRNAs [48]. Since our results demonstrate that C6orf203 also plays an important role in the synthesis of mitochondrial proteins, we investigated their putative interactions with translation-related proteins. Using the Harmonizome collection (http://amp.pharm.mssm.edu/Harmonizome/about#resources), which manages information about genes and proteins from 114 datasets obtained from 66 online resources [49], 260 putative C6orf203 interactors from the *Pathway Commons Protein-Protein Interactions* dataset were extracted. From them, 62 were proteins of the mitoribosome (37 from the large mitoribosome subunit, mt-LSU, and 25 from the small mitoribosome subunit,mt-SSU) (Supp Table 2). In addition, analysis using the NCBI Resources Genes showed 37 putative interactors with C6orf203, 16 of which were MRPLs and 1 was a MRPS (Supp Table 2). Finally, STRING Interaction Network [50] showed 58 proteins that putatively could interact with C6orf203, 15 of them were members of the mitoribosome. Within the top 25 interactions 7 were mitoribosome proteins (Table EV 2).

Then, in order to confirm the possibility that C6orf203 may interact or form part of high molecular weight protein complexes, perhaps the mitochondrial ribosome, we used the cell line HEK293T expressing a TAP-tagged version of C6orf203. Their mitochondria were isolated, solubilized with the relative soft detergent digitonin and subjected to 2D BN/SDS-PAGE followed by Western blot analysis. Our results revealed that C6orf203-TAP was part of several complexes ranging in size from monomeric C6orf203-TAP to high molecular weight complexes. By comparison with the observed supercomplex I+III_2_+IV (SC) signal, as revealed by anti-NDUFS2 antibody, C6orf203 forms part of a supercomplex of approximately 1,500 kDa in size (Fig 9A). These results were confirmed by isokinetic sucrose gradients. This analysis confirmed that C6orf203 forms complexes from low (probably the monomer) to very high molecular weight, being particularly enriched in the fractions corresponding to the 39S mitochondrial ribosome large subunit (mtLSU) (Fig 9B).

**Fig 9.**
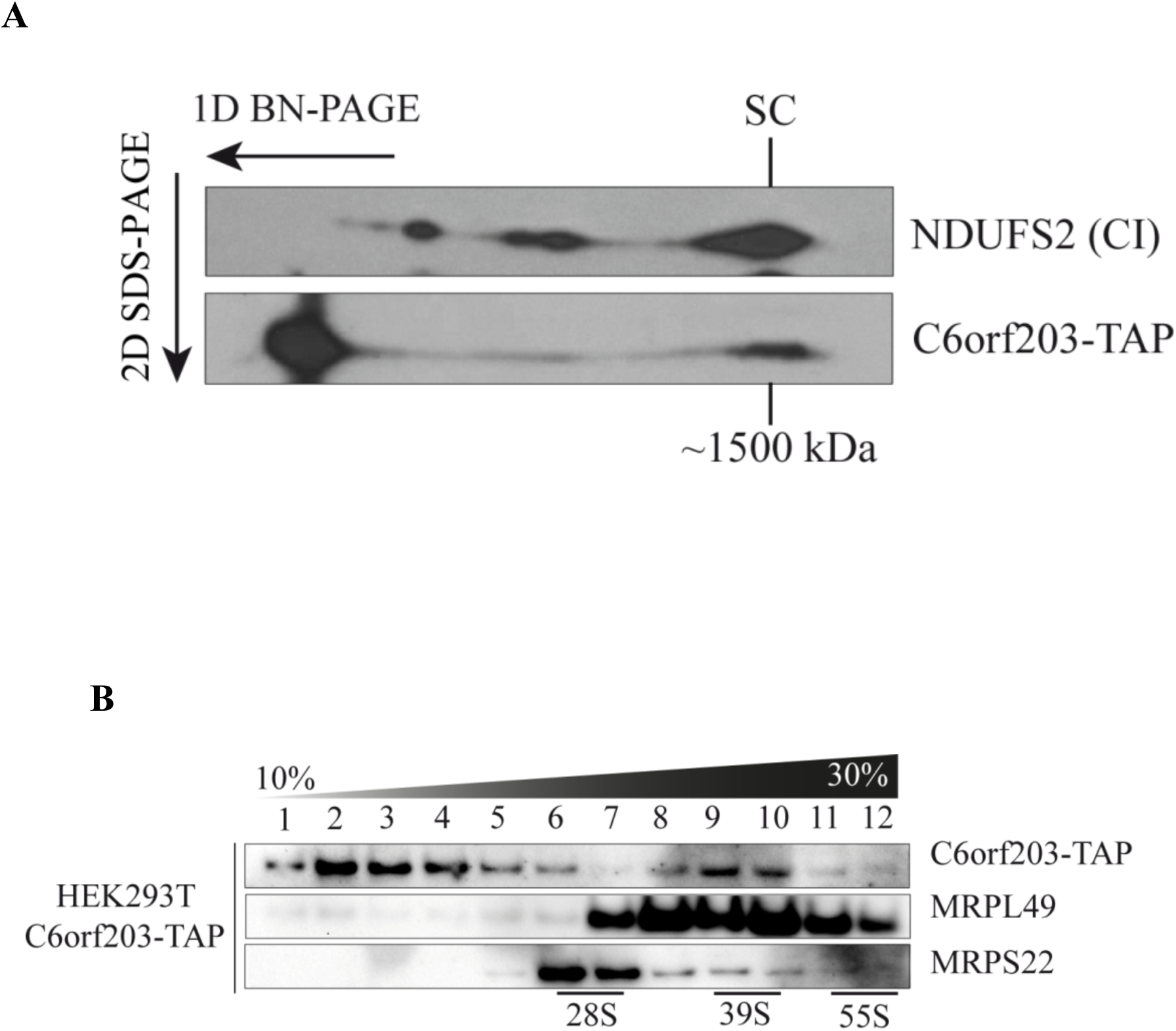
C6orf203 forms high molecular weight complexes. **A** Two dimensional BN-PAGE/SDS-PAGE analysis of mitochondria from HEK293T overexpressing C6orf203-TAP. 80 μg of mitochondrial protein extract were first separated in a BN-PAGE 4%-15% acrilamide and then resolved in a second denaturing dimension (2D-BN/SDS-PAGE). Distribution of C6orf203-TAP was analyzed by Immunoblot using anti-C6orf203. In addition, distribution of NDUFS2, a complex I subunit, was also analyzed as an indicator of complexes molecular weights. The signal of the supercomplex I+III_2_+IV (SC) is shown. **B** Distribution of C6orf203 and mitoribosomal marker proteins on a sucrose density gradient centrifugation. 800 μg of protein from mitochondrial lysates of HEK293T overexpressing C6orf203-Flag were separated through 10%-30% (w/v) isokinetic sucrose gradients, 12 fractions recovered and analyzed by Western blotting using anti-C6orf203, anti-MRPL49 and anti-MRPS22 antibodies. Fractions containing the small subunit (28S), the large subunit (39S) and the monosome (55S) are indicated. n= 2.

### Mitoribosome assembly is not affected by the lack of C6orf203

Our results pointed towards the involvement of C6orf203 in mitochondrial translation being part of high molecular weight complexes that well could correspond to mitochondrial ribosomes. In order to test if C6orf203 was involved in mitoribosomes formation, we analyzed the mitochondrial ribosome biogenesis profile on sucrose gradients followed by Western blot using antibodies recognizing the markers MRPS22, a member of mt-SSU or 28S small subunit, and MRLP49, a subunit of mt-LSU. Thus, we analyzed mitochondria from wild type, C5 and C35 cell lines.

Distribution of MRLP49 and MRPS22 was similar in all cell lines used, strongly suggesting that absence of C6orf203 did not lead to a mitoribosome biogenesis defect (Fig 10). However, it cannot be ruled out it is involved in mitoribosomes function.

**Fig 10.**
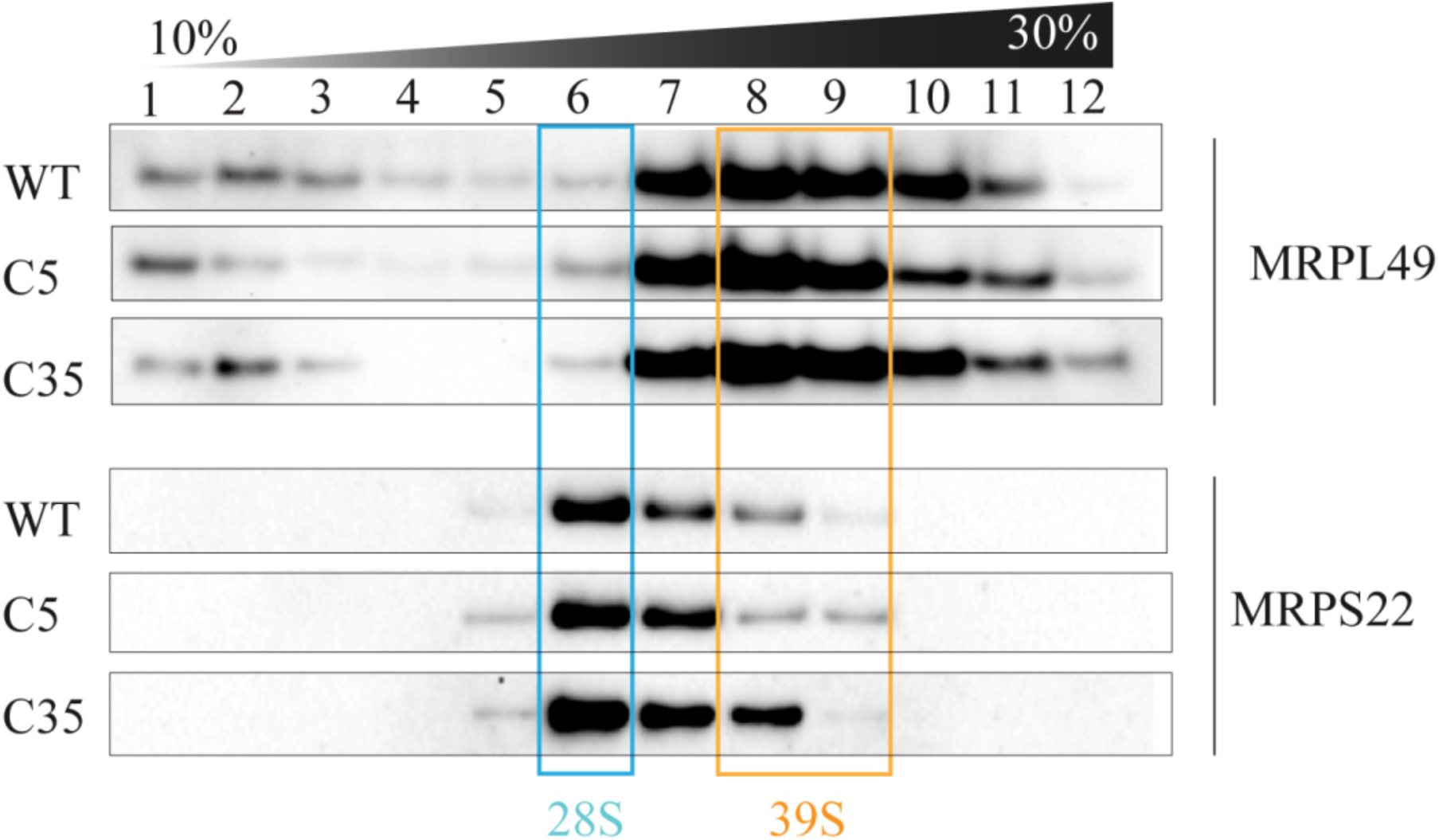
C6orf203 depletion does not affect mitoribosomal assembly. Sedimentation analysis of the large and the small mitoribosomal subunits. 800 ug of mitochondrial extracts prepared from wild type, C5 and C35 cell lines were separated in 10%-30% (w/v) sucrose gradients, fractionated and analyzed by SDS-PAGE and immunoblotting using the indicated antibodies. MRPL49 mark the large ribosomal subunit (39S) and MRPS22 the small ribosomal subunit (28S). n=2.

## Discussion

Mitochondrial OXPHOS function is the result of a coordinated collaboration of hundreds of proteins, of which many are still to be identified and characterized. As such, the identification and characterization of all OXPHOS players is essential to fully understand the complex physiology of this essential biological process. Furthermore, by fully understanding the physiology of the mitochondria we can better comprehend mitochondrial dysfunctions, which result in mitochondrial OXPHOS diseases. Thus, discovering missing OXPHOS-related functional genes opens up the possibility of understanding, diagnosing and of thinking about possible therapeutic approaches for patients with OXPHOS diseases. In our lab, by data mining the *Drosophila* genome and by ortholog analysis, we have identified several genes essential for mitochondrial function, which are coded within bicistronic mRNAs [23] [22, 24], leading us to suggest that this arrangement of mitochondrial genes may be a tendency in this organism (manuscript in preparation). A few weeks ago, Kotrys and collaborators, by a proteomic analysis of the human mitochondria, identified *C6orf203*/*MTRES1* as a new mitochondrial gene [28]. They describe this protein as a factor that, acting at transcriptional level, prevents mitochondrial transcription deficiency under mtDNA depletion induced by BrEt. In our study, we describe the identification of *C6orf203*/*MTRES1* by a different approach, and via its characterization we describe a radical and novel function for *C6orf203*/*MTRES1* in OXPHOS system biogenesis, different than that described by Kotrys et al.

Our analysis of bicistronic mRNAs in *Drosophila* also identified *C6orf203*. In addition, C6orf203 showed a putative efficient mitochondrial targeting sequence (MTS), a conserved RNA binding domain, it was included in MitoCarta, was co-immunoprecipitated with ICT1, an integral component of the human mitoribosome [27] and it was evolutionary conserved. All these evidences made C6orf203 a very interesting gene and worthy of study.

Using the CRISPR/Cas9 genomic edition system in HEK293T cells, an actively respiring cell line, we obtained 6 clones with different edited versions where C6orf203 was not expressed in 5 of them or it was expressed a truncated non-functional version, making these clones the best and easiest model to study and understand the function of C6orf203. Under basal conditions, all *C6orf203*-KO cells had their oxygen consumption and their capacity to grow in galactose as carbon source medium severely affected, a phenotype clearly associated to an alteration of mitochondrial OXPHOS function. Importantly, the ability of a C6orf203-Flagged version to rescue the affected phenotype confirmed the association of the functional defect with the *C6orf203* edition. Thus, these functionally relevant results provided new evidence to support that C6orf203 plays an important role in mitochondrial function under normal growing conditions.

Different approaches revealed that *C6orf203*-KO cells showed OHPHOS global alterations, decreasing: oxygen consumption, mitochondrial ATP synthesis, steady-state levels and the activity of respiratory chain complexes and steady-state levels of mitochondrial encoded OXPHOS subunits. While this general molecular phenotype was compatible with defects in mtDNA decoding, analysis of both mtDNA content and steady-state levels of mitochondrial transcripts under normal growing conditions showed no alterations in the aforementioned processes in cells lacking C6orf203. Moreover, the processing of the selected mtRNAs analyzed were also not affected. In contrast, however, in *C6orf203*-KO cell lines the capacity to synthesize mitochondrial proteins was significantly affected, suggesting that C6orf203 is instead a mitochondrial translation factor, as its physical interaction with the mitochondrial ribosome subunit ICT1 suggested. These results revealed an unexpected new role for C6orf203, being its involvement in translation under normal growing conditions in addition to working as a mitochondrial transcript protection factor. This also reinforced the relevance of using appropriate systems, such as the *C6orf203*-KO cells described, in order to understand the real biological involvement of putative OXPHOS proteins.

Our subcellular fractionation study revealed that C6orf203 behaved as a soluble protein located on the mitochondrial matrix and interacting with IMM. This observation, together with its involvement in mitochondrial translation, its physical interaction with the mitoribosome subunit ICT1 and the well-known interaction of mitoribosome with IMM [51, 52], made us analyze the distribution of C6orf203 in a sediment coefficient-separating sucrose gradient, in comparison with mitoribosome subunits. We confirmed that C6orf203 formed part of very high molecular weight complexes, also confirmed by 2D BN/SDS-PAGE, perhaps the mitoribosome itself. Along these lines, several on-line resources specialized in the analysis of protein-protein interactions, strongly suggested the interaction of C6orf203 with the mitoribosome, showing percentages between 23.8% and 45.9% of mitoribosome proteins as C6orf203 interactors. Since the factors that participate into mitoribosome biogenesis remain largely unknown, we analyzed the putative involvement of C6orf203. The similar kinetics of mitoribosome subunits assembly in wild type cells and in *C6orf203*-KO cells discarded this possibility; however, although the decrease in mitochondrial translation rate in *C6orf203* knockout cells is not due to a defect in mitoribosome assembly, it cannot be ruled out that C6orf203 depletion leads to a serious but not deleterious defect in mitoribosome function. Thus, the specific role of C6orf203 in the synthesis of mitochondrial proteins is still unknown and requires future work. We hypothesize that it could be a member of the mitoribosome with an important but not essential function in the translation process, or it could be a translation activator similar to others IMM-interacting proteins such as Mba1 [45] or PET111 [46]. Furthermore, the phosphorylation of C6orf203 at Ser-106/Ser-110 by a still unknown Kinase [26] suggests that this protein could participate in regulatory events depending on the cellular context.

A final and significant consequence of this work is the possible clinical expression of an important but not complete defect in translation by lack of C6orf203 function. Therefore, patients with a suspected mitochondrial disease caused by C6orf203 mutations probably will not show a deleterious but rather a mild phenotype, unless a synergistic effect with other mutated genes exists. In addition, the apparent double function of C6orf203 as a protector of transcription under mtDNA depletion or as a regulator of translation under normal conditions will surely have different clinical manifestation depending on the degree to which each function is affected. Thus, our study complements, in a timely manner, the conclusions recently derived by Kotrys and collaborators [28].

In conclusion, this work confirms that C6orf203 is a soluble protein located on the mitochondrial matrix, interacting with IMM, which form part of high molecular weight complexes, co-sediments with mitoribosome subunits and is involved in regulating mitochondrial protein synthesis under basal conditions. C6orf203 is therefore a good candidate to be included in clinical panels of genes to identify the origin of some, likely mild, mitochondrial pathologies.

## Materials and methods

### Cell lines, cell culture and transfection experiments

HeLa, HEK293T and Flp-In T-Rex HEK293 cells were cultured in high-glucose DMEM supplemented with 10% FBS, 50 µg/ml uridine, 100 U/ml penicillin and 100 µg/ml streptomycin at 37°C in a humidified 5% CO2 atmosphere. Usually, 8×10^5^ HeLa and HEK293T cells were transfected using Lipofectamine 2000 (Invitrogen) following manufacture’s protocol. Cells stably overexpressing C6orf203-Flag were selected and maintained in DMEM containing 1.5 µg/ml puromycin. Transfection of Flp-In T-Rex HEK293 cells was carried out according to Flp-In™ T-REx™ Core Kit (Invitrogen). Briefly, one day before transfection penicillin/streptomycin were removed from the culture medium. When reached 60-80% confluence, cells were co-transfected with the C6orf203-TAP construct and with plasmid pOG44 using Superfect (Qiagen). After 48h, cells were selected by the addition of blastidicin and hygromycin (Invitrogen). Clones were selected and, when reaching 60-80% confluency, protein expression was induced by using 1 μg/ml doxcycline (Sigma Aldrich) for 24h.

### Constructs

Some constructs were obtained from HEK293T cDNA. Thus, total RNA was previously extracted from cells using the TRIzol® reagent (Sigma) following the manufacturer’s instructions. For cDNA synthesis, the High-Capacity cDNA Reverse Transcription kit (Life Technologies) with 1 μg of total RNA was used.

To construct the C-terminal Flag-tagged version of C6orf203, its cDNA was amplified without the stop codon using the forward primer 5’-CAT***CTTAAG***ATGGCTATGGCTAGTGTTAA-3’ and the reverse primer 5’-TA***GCGGCCGC***TTA<u>CTTGTCGTCATCGTCTTTGTAGTCTTT</u>AGACATTCTCTTCT-3’ that contains the Flag tag (underlined) in frame with C6orf203 coding region and the restriction sites for AflII and NotI (bold and italic), respectively. The PCR product was cloned into the vector pIRESpuro2 using those restriction enzymes (Clontech).

To express C6orf203 fused to eGFP (Enhanced Green fluorescent protein), the C6orf203 coding sequence lacking the stop codon was amplified by PCR using the forward primer 5’-CCA***CTCGAG***ATGGCTATGGCTAGTGTTAAA-3’ and the reverse primer 5’-CCA***GGATCC***TTAGACATTCTCTTCTTAGG-3’ that contain the restriction sites for XhoI and BamHI (bold and italic), respectively. The PCR product was cloned into the corresponding restriction sites of pEGFP-N1 vector (Clontech).

A doxycyclin inducible C6orf203-TAP-tagged (C6orf203-TAP) expression vector was constructed as described in [29]. Final construct was transfected into Flp-In T-Rex HEK293 cells as described above.

Cloning for expression of gRNAs and Cas9 for genome editing is described below. Accuracy of all clones was confirmed by DNA sequencing.

### Generation of C6orf203 deficient cell lines using CRISPR/Cas9 nickase system

For the generation of *C6orf203*-KO cells the CRISPR/Cas9 system based on Cas9-derived nickase and a pair of gRNAs was used [30]. gRNAs were obtained by annealing and cloning complementary primers into vector pX335-U6-Chimeric_BB-CBh-hSpCas9n(D10A) containing humanized *S.pyogenes* Cas9n(D10A) nickase (Addgene, cat. # 42335). Primers were designed to target the exon 2 of *C6orf203* gene using E-CRISP double nickase platform (http://www.e-crisp.org/E-CRISP/) [31]. Thus, the set of primers were Pair 1: 5’-<u>CACC</u>**GCTCACTCATCTCATCATGA-3’** and 5’-<u>AAAC</u>TCATGATGAGATGAGTGAGC-3’; Pair 2: <u>CACC</u>**GTATGATGTTGTCCTGAAGA** and <u>AAAC</u>TCTTCAGGACAACATCATAC).

*Bbs*I restriction site overhang ends are underlined. gRNAs sequences to be expressed are in bold. Annealed corresponding pairs of primers were cloned into the *Bbs*I cloning sites of vector pX335-U6-Chimeric_BB-CBh-hSpCas9n(D10A) by standard procedures using *E.coli DH5α*. Accuracy of all clones was confirmed by DNA sequencing. Before cloning, we PCR amplified and sequenced the genomic region of HEK293T cells to be edited in order to confirm that there were no differences in this region with published database sequences. Primers used for that were Fwd: 5’-TTCTCACTGAGACTCCCAGG-3’ and Rev: 5’-TGGGCCAAGTGATTTGTAATGC-3’). Thus, gRNAs expressing plasmids were transfected in HEK293T cells. 24 hours after transfection, edition of *C6orf203* was verified in the pool of cells by genomic DNA extraction and PCR using the same primers. Next, 40 individual clones were isolated and C6orf203 expression analyzed by Western Blot with C6orf203 antibody (Abcam, ab151066). Then, edition of those lacking C6orf203 expression was confirmed by PCR and sequencing.

### Subcellular localization of C6orf203

To analyze the subcellular localization of C6orf203, HEK293T or Hela cells were transiently transfected with the peGFP-N1 derivative construct that expresses C6orf203 fused to eGFP at its C-terminal. 24 hours after transfection cells were incubated with medium containing 100 nM MitoTracker Red CMXRos (Thermo Scientific), a mitochondrion-selective dye, for 30 min at 37°C. Cells were washed twice with PBS and fixed with 4% PFA for 20 min. After three washes with PBS, images were obtained in a confocal microscope with a 63x oil objective (Carl Zeiss LSM 700, Oberkochen, Germany).

### Mitochondrial subfractionation, proteinase K protection assay and carbonate extraction

An enriched mitochondria fraction was previously isolated from 4×10^7^ HEK293T cells by differential centrifugations following the protocol described in [32]. Then, the mitochondrial fraction was purified by ultracentrifugation at 100,000 g for 2 hours in a discontinuous sucrose gradient (five fractions from 1.6 M to 0,8 M sucrose). Purified mitochondria were obtained from the sucrose 1.4/1.2 M interphase and washed.

Proteinase K protection assay was performed as described in [33] with slight modifications. 200 μg of purified mitochondria were incubated with 100 μg/ml proteinase K in the presence or absent of a swelling buffer (2mM HEPES pH 7.4) for 30 minutes on ice. The reaction was stopped by addition of Protease Inhibitor Cocktail (Sigma, Saint Louis, MO, USA). Proteins were then precipitated with 15% trichloroacetic acid (TCA) at 4°C for 30 min and centrifuged at 20,000g, washed with cold acetone and resuspended in Tris buffer previous to immunoblotting.

For carbonate (Na_2_CO_3_) extraction treatment, 200 μg of purified mitochondria were resuspended in SEH buffer (20mM HEPES-KOH pH 7.4, 0.6M sorbitol, 2mM MgCl2, 1mM EGTA pH 8.0 and 1X Protease Inhibitor Cocktail) containing 0.1M Na_2_CO_3_ pH 11 and incubated for 30 min at 4°C. Soluble and insoluble fractions were obtained by centrifugation at 20,000 g for 30 min at 4°C. Membrane fraction (pellet) was resuspended in 40 µl of 1X Laemmli loading buffer. Soluble proteins (supernatant) were precipitated using 15% TCA and resuspended in 40 µl of 1X Laemmli loading buffer. Both samples were resolved on a 12% SDS-PAGE gel and analyzed by immunoblotting.

### Growth curves

Cell growth was determined by seeding 2.5×10^4^ cells/well of wild-type HEK293T, *C6orf203*-KO or *C6orf203*-KO cells expressing the flag-tagged version of C6orf203 in poly-lysine treated 6-well plates. Growth in DMEM medium containing either 4.5g/l glucose or 4,5 g/l galactose as a carbon source was measured over 4 days. Cells were trypsinized and counted (in duplicate for each condition) every 24 hours. Three different growth curves in independent experiments were carried out.

### Measurement of oxygen consumption

Wild-type HEK293T cells, *C6orf203*-KO cells and *C6orf203*-KO cells expressing the flag-tagged version of C6orf203 were grown in DMEM medium containing 0.9 g/l galactose for 12 hours. Intact cells were then collected and their basal respiration was measured in DMEM-galactose using a Clark-type oxygen electrode (Hansatech Instruments) as described [34]. Assays were performed in three independent experiments.

### ATP measurement

Measurement of mitochondrial steady-state ATP levels was carried out in cells incubated with 5 mM 2-deoxy-D-glucose/1 mM pyruvate and no carbon source using the luminometric luciferin-luciferase-based method (CLS II, Roche Applied Science) as described in [35]. Experiments were performed in duplicate on at least three independent days.

### Enzymatic activity of OXPHOS complexes

The activity of the respiratory chain complexes and the enzyme citrate synthase was measured as previously described [36] using a Beckman Coulter DU 800 spectrophotometer.

### Pulse labeling of mitochondrial translation products

Synthesis of mtDNA encoded proteins was analyzed by pulse labeling experiments according to [37] with some modifications. Briefly, exponentially growing 4×10^5^ HEK293T cells were cultured 90 min with 200 µCi/ml [S^35^]-methionine (Perkin Elmer) in methionine and cysteine-free DMEM containing 100 µg/ml of emetine, an inhibitor of cytosolic translation. Then, cells were harvested and lysed in RIPA lysis buffer [150mM NaCl, 1% Nonidiet P40, 0.1% SDS, 0.5% deoxycholate, 1mM EDTA and 50mM Tris-HCl (pH 8.0) with Protease Inhibitor Cocktail (Roche)] and incubated for 20 min on ice. Insoluble material was removed by centrifugation at 14,000g for 5 minutes at 4°C. Protein concentration was determined using Pierce BCA Protein Assay kit (Thermo Scientific) and 50 µg of each protein sample were resolved onto 17.5% SDS-PAGE gels. After electrophoresis, gels were fixed, incubated in Amplify Fluorographic Reagent (GE Healthcare, ref. NAMP100) for 15 min and dried. Labeled mitochondrial proteins were visualized through direct autoradiography. Gels were finally stained with Coomassie brilliant blue to confirm equal loading.

### DNA isolation and quantification of mtDNA

Total DNA was isolated from cultured cells using a commercial kit (Qiagen, UK), following the manufacturer’s recommendations. The mtDNA content in 50 ng of total DNA was measured by SYBR green real-time polymerase chain reaction (PCR) using the Applied Biosystems Step-One Plus real-time thermocycler according to [24] consisting of an initial 10 min denaturation step at 95°C followed by 40 cycles of denaturation (15 sec at 95°C) and annealing/extension (1min at 60°C). The analyzed genes were the mitochondrial *12S-rRNA* gene (or *MT-RNR1*) and the nuclear *HPRT1* gene. Pairs of primers used for *12S-rRNA* were forward 5’-CCACGGGAAACAGCAGTGAT-3’ and reverse 5’-CTATTGACTTGGGTTAATCGTGTGA-3’ and for *HPRT1* were forward 5’-CCTGGCGTCGTGATTAGTGAT-3’ and reverse 5’-AGACGTTCAGTCCTGTCCATA-3’. For determining mtDNA copy number, an independent standard curve was generated for each gene (*12S-rRNA* and *HPRT1*). mtDNA copy number values were expressed by the ratio *12S rRNA/HPRT1*.

### mtRNA content and intermediates

Total RNA was extracted and cDNA synthesized as in “Constructs” section followed by qPCR using conditions as detailed in the previous point.

Genes analyzed and the primers used for it were: *12S-rRNA* (primers were the same than those used for mtDNA quantification), *MT-CO1* (FW: 5’-CTCTTCGTCTGATCCGTCCT-3’ and REV: 5’-ATTCCGAAGCCTGGTAGGAT-3’) *MT-CO2* (FW: 5’-CGCCCTCCCATCCCTACGCA-3’ and REV: 5’-CCGCCGTAGTCGGTGTACTCG-3’) *MT-*ND1 (FW: 5’-GCACTGCGAGCAGTAGCCCA-3’ and REV: 5’-TGGCCAAGGGTCATGATGGCA-3’) and *MT-ND6* (FW: 5’-GATTGTTAGCGGTGTGGTCGGGT-3’ and REV: 5’-GACCTCAACCCCTGACCCCCA-3’).

In order to analyze mtRNAs intermediates, and like an additional approach for mtRNA quantification, northern blot assays were carried out according to [38]. Briefly, 5 μg of total RNA from wild-type HEK293T and *C6orf203*-KO cells were resolved on a 1.5% agarose gel containing 18% formaldehyde and transferred to a ZETA-PROBE GT membrane (Bio-Rad). Probes used for hybridization were ^32^P-labeled specific antisense ssRNA corresponding to *16S-rRNA* gene (or *MT-RNR2*), *MT-CO1* and *MT-ATP8/6* genes. For riboprobes synthesis, PCR using reverse primers including the T7 phage promoter sequence at their 5’end and standard forward primers were used: 16S-rRNA_FW: 5’-ACCCAAATAAAGTATAGGCGATAGAAATTGAAACCTGGCGCAATAG-3’, 16S-rRNA_REV: 5’-<u>CGGTAATACGACTCACTATAGGGAGA</u>TCCTAGTGTCCAAAGAGCTG-3’; MT-CO1_ FW: 5’-CCTACTCCTGCTCGCATCTG-3’, MT-CO1_ REV: 5’-<u>CGGTAATACGACTCACTATAGGGAGA</u>CGGCGGGGTCGAAGAAGGTG-3’ and ATP8/6_FW: 5’-ATGCCCCAACTAAATACTACC-3’, ATP8/6_R: 5’-<u>CGGTAATACGACTCACTATAGGGAGA</u>TTGGGTGGTTGGTGTAAATGA-3’. T7 phage promoter is underlined. 200ng of the corresponding PCR products were used as templates for transcription using the MEGAscript T7 transcription kit (Ambion) in presence of 30μCi of [α-^32^P] UTP following the manufacte’s instructions. The 16S-rRNA gene, MT-CO1 and MT-ATP8/6 riboprobes were 399 nts., 343 nts. and 450 nts in size respectively. Gels were transferred to a Zeta-Probe membrane (Bio-Rad) that was subjected to a 1200×100 μJ/cm^2^. Later, the membranes were incubated with PerfectHyb Plus Hybridization Buffer (Sigma) for 2 hours at 60°C before addition of denatured riboprobes. The riboprobes were incubated at 60°C overnight and the membranes were washed and developed using x-ray films.

### Immunoblotting (Western Blot)

Cellular or mitochondrial pellets were resuspended in RIPA lysis buffer and protein extracts obtained as above. Equal amounts of protein (50 to 100 µg) were subjected to denaturing SDS-PAGE (regularly 12% polyacrylamide gels) resolution and then transferred to PVDF membranes (Immobilon-FL, Millipore). After blocking 1h-3h at room temperature with 5% (w/v) non-fat dry milk in TBS with 0.1 % (v/v) Tween-20, membranes were incubated with specific primary antibodies (see below). The blots were finally incubated with the corresponding horseradish peroxidase-conjugated secondary antibodies and visualized using ECL. Protein bands were quantified using Image J. The following primary antibodies were used in this work (ordered by appearance in results): αTubulin (Proteintech, HRP-66031), TOM20 (Abcam, ab56783), OPA1 (Abcam, ab42364), MnSOD (Merck Millipore, 06-984), C6orf203 (Abcam, ab151066), MT-CO1 (Abcam, ab14705; OXPHOS system Complex IV), SDHA (459200; TermoFisher), VDAC (Abcam, ab15895), COX5A (Abcam ab110262; OXPHOS system Complex IV), UQCRC2 (Abcam, ab14745; OXPHOS system Complex III), NDUFA9 (Abcam ab14713; OXPHOS system Complex I), ATP5A1 (Abcam, ab14748; OXPHOS system Complex IV), SDHB (Abcam, ab14714; OXPHOS system Complex II), NDUFB8 (Abcam, ab110242; OXPHOS system Complex I), MT-ND1 (a kind gift of Dr. Anne Lombès; OXPHOS system Complex I), MT-CO2 (Abcam, ab110258; OXPHOS system Complex IV), MRPL49 (Proteintech, 15542-1-AP), MRPS22 (Proteintech, 10984-AP), NDUFS2 (Abcam, ab110249; OXPHOS system Complex I).

### Blue Native-PAGE and Two-dimensional BN/SDS-PAGE

Blue Native PAGE (BN-PAGE) and two-dimensional BN/SDS-PAGE experiments were performed as previously described [39].

### Sucrose gradient analysis of mitoribosomes

Mitoribosomes were purified on sucrose gradients as previously described [40]. Briefly, 2.4×10^7^ HEK293T cells were harvested in PBS containing 10 mM MgCl_2_, centrifuged 5 min at 300g, incubated for 15 min at 4°C in lysis buffer (50 mM Tris-HCl pH 7.2, 10 mM Mg(CH3COO)2, 40 mM NH4Cl2, 100 mM KCl, 1mM PMSF and 1% n-dodecyl-β-D-maltoside) and centrifugated at 14,000g for 20 min at 4°C. Then, 800 µg of protein from the supernatant (in a final volume of 200 µl) was loaded onto 4 ml of a 10% −30% (w/v) linear sucrose gradient and centrifuged at 74,000g at 4°C for 6 h (Beckman MLS-50 rotor). Fractions (250 µl) were collected from the top of the tube and analyzed by Western blotting. The distribution of mitoribosomal subunits through the gradient was analyzed using MRPL49 and MRPS22 antibodies.

### Statistical Analyses

All results are presented as means ± standard deviation of the mean (SEM) of at least three independent experiments. Statistical significance was assessed by using two-tailed unpaired Student’s t test. A p-value < 0.05 (*p < 0.05, **p < 0.01, and ***p < 0.001) was considered statistically significant.

## Acknowledgements

This work was supported by Instituto de Salud Carlos III, http://www.isciii.es/, PI16/00789 to R.G. and M.A.F-M., FEDER funds from the E.U. to R.G. and funds from Raregenomics-CM consortium, https://www.rare-genomics.com/, S2017/BMD-3721 to R.G. and M.A.F-M.

We are grateful to Bruno Sainz Jr. for helpful comments on the manuscript.

We want to make a special remembrance to Leo Nijtmans, author of this work, a great scientist and a great person that recently passed away.

## Author contributions

RG and MAF-M conceived the project, MAF-M designed the experiments and wrote the core of the manuscript, SP-Z performed the majority of the experiments and participated in writing the manuscript and making the figures, LV-F performed the submitochondrial localization, ATP synthesis, northern blots experiments and participated in writing the manuscript, CG-P carried out western blots and oxygen consumption measurements, LM supported some experimental work, SP-Z and LS-C supervised by LN made the C6orf203-TAP construct and its analysis.

## EXPANDED VIEW

**Fig EV 1.**
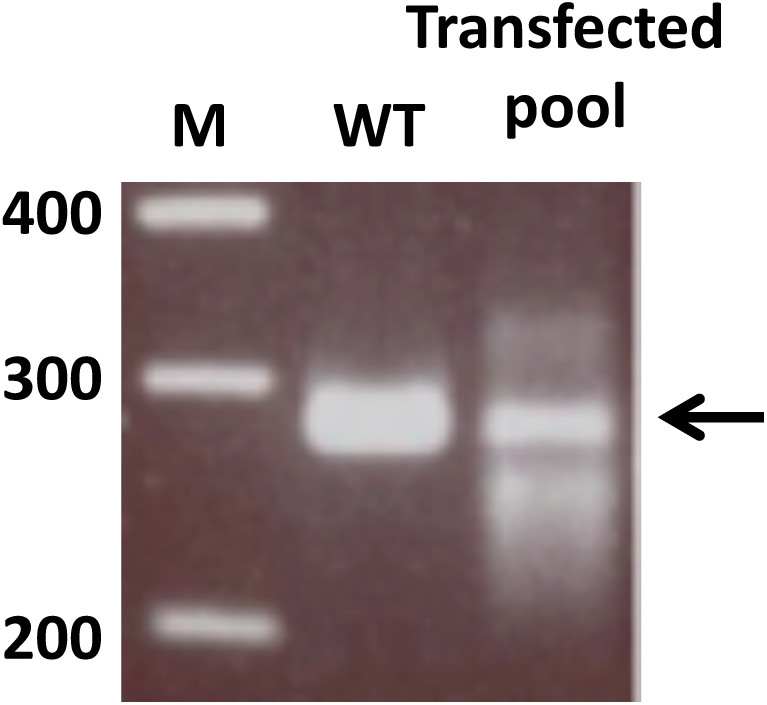
Edition of C6orf203 within the pool of CRISPRized HEK293T cells. 24 hours after transfection of 8×10^5^ HEK293T cells with plasmid expressing gRNAs and CRISPR/Cas9n(D10A) nickase, cells were harvested and the genomic DNA of 10^5^ of them was extracted and analyzed by PCR, amplifying a 299 bp in size including the edited region as described in Materials and Method. Multiple variations in the PCR product size were observed. Densitometry indicates that the intensity of the 299 bp band from the pool edited cells approximately corresponds with the 36% of the intensity of the band from non-edited cells.

**Fig EV 2.**
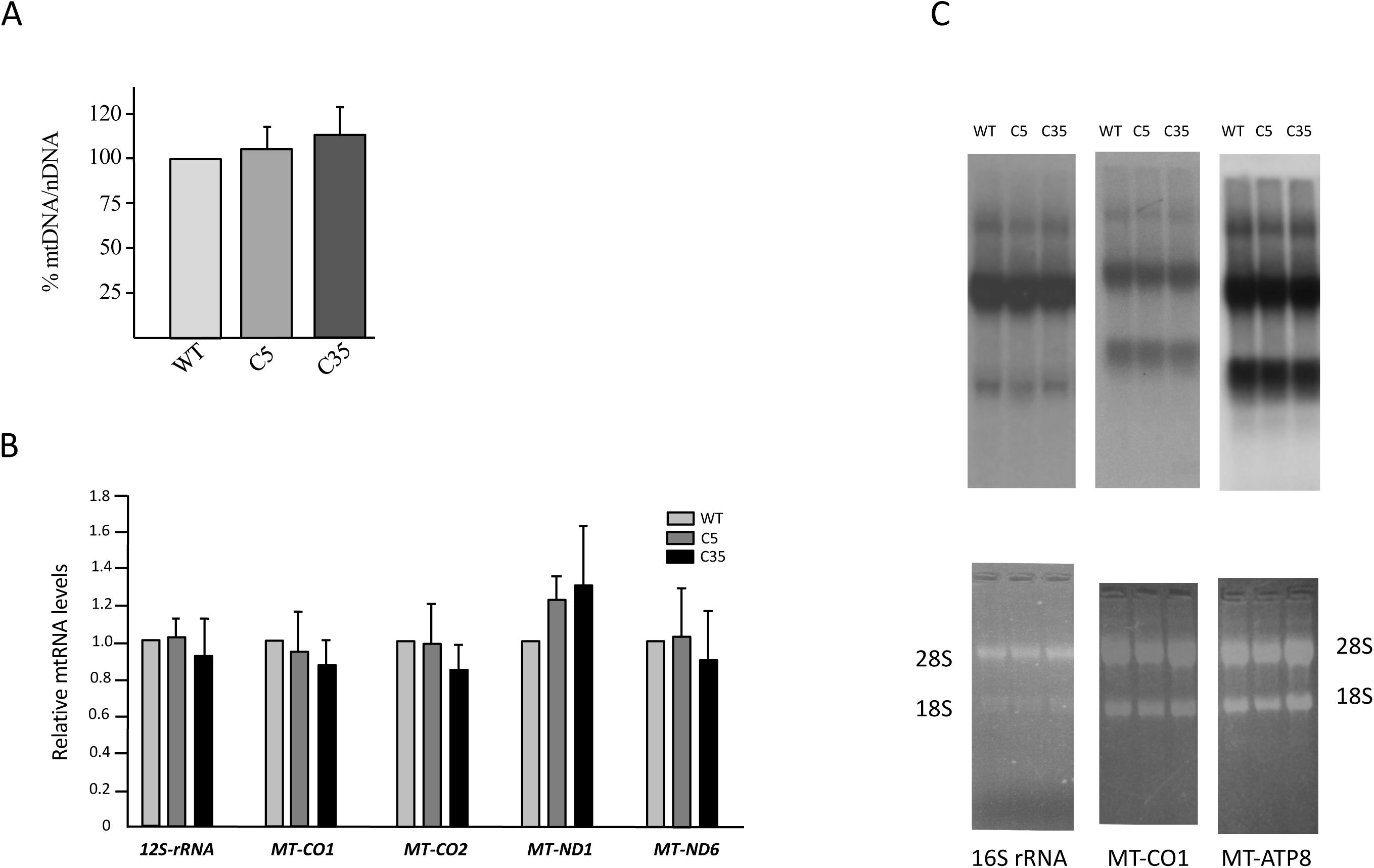
Analysis of mtDNA, mtRNA steady-state levels and mtRNA processing. **A** For quantification of mtDNA content a real-time PCR was performed. Total DNA from wild type and *C6orf203*-KO cell lines was used as template. mtDNA was quantified using specific primers for the *12S-rRNA* mitochondrial gene using the levels of the nuclear encoded gene *HPRT1* for normalization. No significant differences were seen. **B** The relative amount of specific mtRNAs was determined by RT-qPCR in exponentially growing wild type, C5 and C35 cells. The mtRNA content was determined using specific primers for *12S-rRNA*, *MT-CO1, MT-CO2, MT-*ND1 and *MT-ND6* genes. Data represented are normalize to mRNA levels of the nuclear-encode control gene *HPRT1*. No significant differences were seen. **C** Northern blot assays were carried out using 5 μg of total RNA from wild type, C5 and C35 cells run in agarose gel containing 18% formaldehyde (lower panels; loading controls) and transferred to ZETA-PROBE GT membrane (Bio-Rad). Probes used for hybridization were ^32^P-labeled specific antisense ssRNAs corresponding to *16S-rRNA* gene (or *MT-RNR2*), *MT-CO1* and *MT-ATP8/6* genes. The riboprobes were incubated at 60°C overnight and the membranes were washed and developed using x-ray films (upper panels). No significant differences were seen in the mtRNAs forms. Since the gels are from different experiments the scale is not the same and it is not possible to superimpose the images.

**Table EV 1.** Edited alleles of six different C6orf203-KO clones. Sequencing of edited alleles of C6orf203-KO clones revealed variations such as deletion (△), frameshift (fs) and stop codon (*). Since HEK293T is a hypotriploid cell line [53] in some cases three different edited alleles are shown. Nomenclature is according the Human Genome Variation Society and changes are referred to ENST00000405204.6 (DNA) and Q9P0P8 (protein).

| Clone | DNA | Protein (*in silico*) |
| --- | --- | --- |
| **1** | c.327_448del (Δ122pb) | p.(Glu109Aspfs*18) |
| | c.426_462del (Δ37pb) | p.(Phe142Leufs*17) |
| **2** | c.327_448del (Δ122pb) | p.(Glu109Aspfs*18) |
| **5** | c.336_446del(Δ111pb) | p.(His112_Lys149del) |
| | c.414_455del (Δ41pb) | p.(Val139_Leu152del) |
| **9** | c.339_445del(Δ107pb) | p.(Asp113Glufs*18) |
| | c.416_457del (Δ42pb) | p.(Val139_Leu152del) |
| | c.421_447del (Δ27pb) | p.(Ser141_Lys149del) |
| **11** | c.421_452del (Δ32pb) | p.(Ser141Alafs*16) |
| **35** | c. 329_446del(Δ118pb) | p.(Ser110Argfs*4) |
| | c. 336_446del(Δ111pb) | p.(His112_Leu149del) |
| | c.344_448del(Δ105pb) | p.(Met115_Lys149del) |

**Table EV 2.**
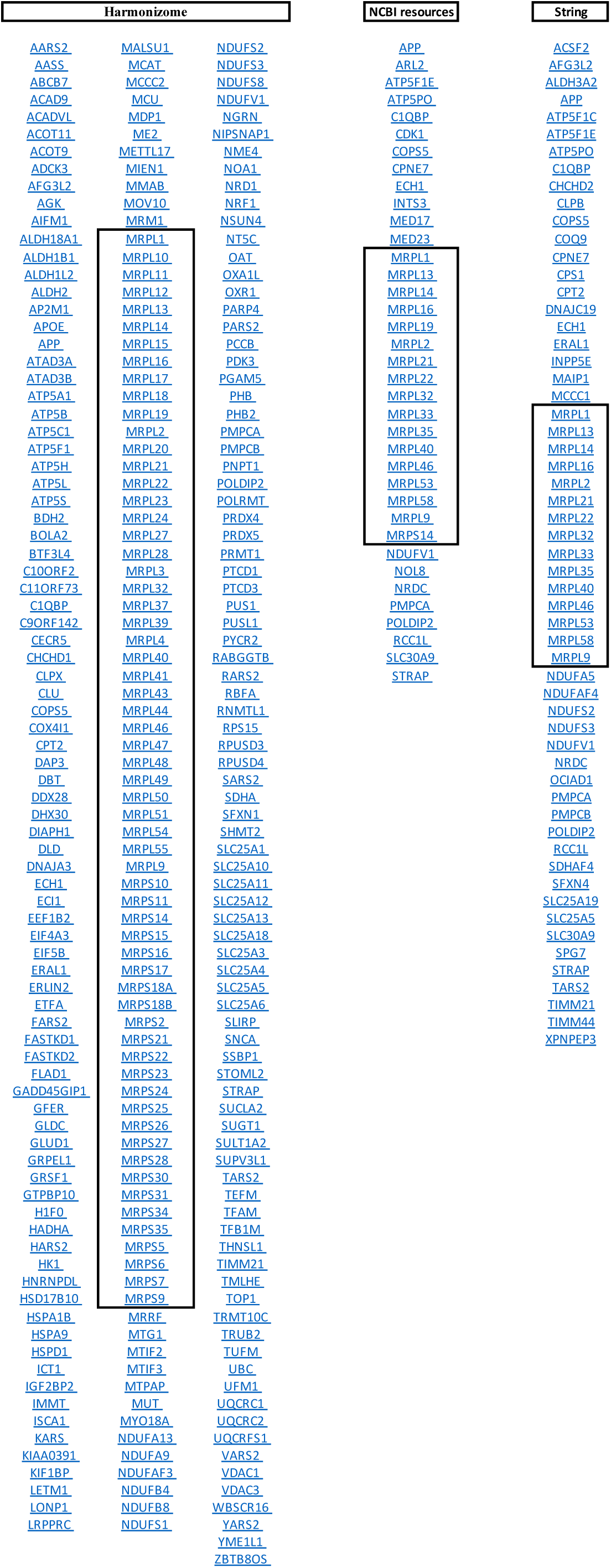
C6orf203 putative interactors. The interactors assigned to c6orf203 by three different resources are shown in an Excel file. The Harmonizome collection, managing proteins from 114 datasets, shows 260 proteins (in three columns) candidate to interact with C6orf203 from which 23,8% are members of mitoribosome (squared). The NCBI resources assign to 37 proteins the possibility to interact with C6orf203, belonging 45,9% to the mitochondrial ribosome (squared). In its turn, STRING Interaction Network shows 58 proteins with different evidences of interaction with C6orf203. From the later, 25,8% are mitochondrial ribosome proteins (squared).

